# Evaluation of IL-1 blockade as an adjunct to linezolid therapy for tuberculosis in mice and macaques

**DOI:** 10.1101/792390

**Authors:** Caylin G. Winchell, Bibhuti B. Mishra, Jia Yao Phuah, Mohd Saqib, Samantha J. Nelson, Pauline Maiello, Chelsea M. Causgrove, Cassaundra L. Ameel, Brianne Stein, H. Jacob Borish, Alexander G. White, Edwin C. Klein, Matthew D. Zimmerman, Véronique Dartois, Philana Ling Lin, Christopher M. Sassetti, JoAnne L. Flynn

## Abstract

In 2017 over 550,000 estimated new cases of multi-drug/rifampicin resistant tuberculosis (MDR/RR-TB) occurred, emphasizing a need for new treatment strategies. Linezolid (LZD) is a potent antibiotic for drug-resistant Gram-positive infections and is an effective treatment for TB. However, extended LZD use can lead to LZD-associated host toxicities, most commonly bone marrow suppression. LZD toxicities may be mediated by IL-1, an inflammatory pathway important for early immunity during *M. tuberculosis* infection. However, IL-1 can contribute to pathology and disease severity late in TB progression. Since IL-1 may contribute to LZD toxicity and does influence TB pathology, we targeted this pathway with a potential host-directed therapy (HDT). We hypothesized LZD efficacy could be enhanced by modulation of IL-1 pathway to reduce bone marrow toxicity and TB associated-inflammation. We used two animal models of TB to test our hypothesis, a TB-susceptible mouse model and clinically relevant cynomolgus macaques. Antagonizing IL-1 in mice with established infection reduced lung neutrophil numbers and partially restored the erythroid progenitor populations that are depleted by LZD. In macaques, we found no conclusive evidence of bone marrow suppression associated with LZD, indicating our treatment time may have been short enough to avoid the toxicities observed in humans. Though treatment was only 4 weeks (the FDA approved regimen at the time of study), we observed sterilization of the majority of granulomas regardless of co-administration of the FDA-approved IL-1 receptor antagonist (IL-1Rn), also known as Anakinra. However total lung inflammation was significantly reduced in macaques treated with IL-1Rn and LZD compared to LZD alone. Importantly, IL-1Rn administration did not impair the host response against Mtb or LZD efficacy in either animal model. Together, our data support that inhibition of IL-1 in combination with LZD has potential to be an effective HDT for TB and the need for further research in this area.

## Introduction

Tuberculosis (TB) remains the top cause of death by a single infectious agent, with an estimated 10 million new cases of active TB and 1.3 million deaths in 2017 alone (1). Antibiotic treatment regimens are long and multi-drug resistant (MDR) and extensive-drug resistant (XDR) *Mycobacterium tuberculosis* (Mtb) strains have emerged, complicating treatment. Even those patients that are cured of the infection can suffer permanent deficits in lung function that result from inflammation and fibrosis (2). Host-directed therapies (HDTs) have been proposed as a potential option for improving therapy. Depending on the strategy, HDTs can function to enhance antimicrobial immune responses and shorten therapy, or inhibit pathological inflammation (3). Since HDTs would be used as part of a multi-drug regimen, targeting mechanisms that increase drug exposure or decrease toxicity are also possible. While some HDT strategies hold promise, very few have been rigorously tested in pre-clinical models (4).

Interleukin-1 (IL-1) has been implicated in TB disease severity and inflammation, making it a possible target of HDT. This cytokine plays an important yet complicated role in TB disease progression. The susceptibility of mice lacking critical mediators of IL-1 signaling indicates that initial production of IL-1 upon Mtb infection is essential for establishing protective immune responses necessary for disease control (5–8). In contrast, IL-1 production is regulated after the onset of adaptive immunity, via multiple mechanisms including IFNγ production(8), which acts via the induction of nitric oxide synthase 2 (NOS2)-dependent nitric oxide to inhibit IL-1β processing(9). Persistent IL-1 signaling can contribute to the accumulation of disease-promoting neutrophils in susceptible mice, and genetic variants that result in higher IL-1β production are associated with increased disease severity and neutrophil accumulation in humans (9–11). Given that HDT is designed to be administered to chronically infected patients during treatment when persistent IL-1 production can play a pathological role, it could be beneficial to block the inflammation and disease promoting activities of this cytokine.

IL-1 may also play a role in the toxicity of linezolid (LZD), an increasingly important antibiotic for the treatment of drug-resistant TB, highlighted by its recent inclusion in a newly approved therapy for MDR-TB (12). While LZD has shown efficacy against XDR and MDR-TB, its wide-spread use has been limited by severe host toxicities that occur after more than 4 weeks of treatment (13, 14). Over the 6-20 month treatment course necessary to treat resistant TB, both reversible bone marrow suppression and irreversible neuropathies are common clinical manifestations (15). LZD-associated toxicities are generally attributed to the inhibition of mitochondrial translation and LZD-mediated bone marrow suppression is promoted by the subsequent mitochondrial damage. This damage acts on the NOD-like receptor family, pyrin domain containing 3 (NLRP3) protein that has been shown to be necessary for LZD-mediated bone marrow suppression in mice (16). NLRP3 forms an inflammasome complex containing caspase-1, which cleaves a number of substrates resulting in cell death and/or the release of active of IL-1β. While the importance of NLRP3 in bone marrow suppression is clear, the relative roles of inflammasome activation and IL-1 signaling remain uncertain.

Based on these studies, inhibiting the IL-1 pathway as a potential HDT could serve two purposes: first, to alleviate LZD-associated host toxicity and second, to reduce the pathology associated with unchecked IL-1 signaling during TB disease. Due to the pro-inflammatory nature of the IL-1 pathway, strict regulatory mechanisms exist within the host to quell this pathway. IL-1 receptor antagonist (IL-1Rn) is a protein produced constitutively at low levels that can increase in response to a variety of cytokine signals. IL-1Rn serves as a decoy ligand for the IL-1receptor type 1 (IL-1R1), blocking signal transduction and subduing activation of downstream pro-inflammatory pathways (17). Anakinra is an FDA-approved recombinant IL-1Rn that is used to treat rheumatoid arthritis and other inflammatory disorders. As there are no FDA approved drugs to inhibit inflammasome activation, inhibition of the IL-1 pathway with biologics like Anakinra is currently the only feasible strategy to modulate this pathway in humans (18).

We hypothesized that the combination of Anakinra (herein referred to as IL-1Rn) with LZD for treatment of active TB disease would reduce LZD-associated toxicities and host inflammation. While there is little rationale to expect IL-1 blockade to enhance bacterial clearance by the antibiotic, suppression of both inflammation and LZD toxicity could provide a significant benefit. To test this concept, we employed two established TB animal models to assess differing aspects of host responses to LZD and IL-1R1 blockade. We used multiple strains of mice to model distinct disease states and dissect the relative importance of the inflammasome and IL-1 signaling in evaluating HDT efficacy. As a translational model, we used cynomolgus macaques in combination with [^18^F] FDG PET CT serial imaging to track TB disease progression, including inflammation, before and during drug regimens (19). Cynomolgus macaques present a similar spectrum of Mtb infection as humans with pathology, granuloma structure and diversity, that recapitulates human TB (20–22). Importantly, we designed our study in macaques to reflect clinical standards, adhering to FDA guidelines for dosage and time frame of LZD and IL-1Rn administration. Together, our data indicate that IL-1 blockade alleviates LZD-mediated bone marrow (BM) suppression in mice and may accelerate the resolution of inflammation in both mice and macaques with TB.

## Materials and Methods

### Ethics Statement

All experimental manipulations, protocols, and care of the animals were approved by the University of Pittsburgh School of Medicine Institutional Animal Care and Use Committee (IACUC). The protocol assurance number for our IACUC is A3187-01. Our specific protocol approval numbers for this project are 15117082, 16017370, 18124275, 13011368 and 16027525. The IACUC adheres to national guidelines established in the Animal Welfare Act (7 U.S.C. Sections 2131 - 2159) and the Guide for the Care and Use of Laboratory Animals (8^th^ Edition) as mandated by the U.S. Public Health Service Policy.

All macaques used in this study were housed at the University of Pittsburgh in rooms with autonomously controlled temperature, humidity, and lighting. Animals were singly housed in caging at least 2 square meters that allowed visual and tactile contact with neighboring conspecifics. The macaques were fed twice daily with biscuits formulated for nonhuman primates, supplemented at least 4 days/week with large pieces of fresh fruits or vegetables. Animals had access to water *ad libitem*. Because our macaques were singly housed due to the infectious nature of these studies, an enhanced enrichment plan was designed and overseen by our nonhuman primate enrichment specialist. This plan has three components. First, species-specific behaviors are encouraged. All animals have access to toys and other manipulata, some of which will be filled with food treats (e.g. frozen fruit, peanut butter, etc.). These are rotated on a regular basis. Puzzle feeders foraging boards, and cardboard tubes containing small food items also are placed in the cage to stimulate foraging behaviors. Adjustable mirrors accessible to the animals stimulate interaction between animals. Second, routine interaction between humans and macaques are encouraged. These interactions occur daily and consist mainly of small food objects offered as enrichment and adhere to established safety protocols. Animal caretakers are encouraged to interact with the animals (by talking or with facial expressions) while performing tasks in the housing area. Routine procedures (e.g. feeding, cage cleaning, etc) are done on a strict schedule to allow the animals to acclimate to a routine daily schedule. Third, all macaques are provided with a variety of visual and auditory stimulation. Housing areas contain either radios or TV/video equipment that play cartoons or other formats designed for children for at least 3 hours each day. The videos and radios are rotated between animal rooms so that the same enrichment is not played repetitively for the same group of animals.

All animals are checked at least twice daily to assess appetite, attitude, activity level, hydration status, etc. Following *M. tuberculosis* infection, the animals are monitored closely for evidence of disease (e.g., anorexia, weight loss, tachypnea, dyspnea, coughing). Physical exams, including weights, are performed on a regular basis. Animals are sedated prior to all veterinary procedures (e.g. blood draws, etc.) using ketamine or other approved drugs. Regular PET/CT imaging is conducted on most of our macaques following infection and has proved very useful for monitoring disease progression. Our veterinary technicians monitor animals especially closely for any signs of pain or distress. If any are noted, appropriate supportive care (e.g. dietary supplementation, rehydration) and clinical treatments (analgesics) are given. Any animal considered to have advanced disease or intractable pain or distress from any cause is sedated with ketamine and then humanely euthanatized using sodium pentobarbital.

### Mice, infection and treatment

C57BL/6 (stock no. 000664), *Nos2^-/-^* (B6.129P2-Nos2^tm1Lau^/J, stock no. 002609), C3HeB/FeJ (stock no. 00658), and *Nlrp3^-/-^* (B6.129S6-*Nlrp3^tm1Bhk^*/J, stock no. 021302) were purchased from the Jackson Laboratory. Breeding pairs of Caspase-1/11 double knock out mice were kindly provided by Prof. Katherine Fitzgerald of the Department of Infectious Diseases at University of Massachusetts (UMASS) Medical School and bred in house. Mice were housed under specific pathogen-free conditions, and in accordance with the UMASS Medical School, IACUC guidelines. All mouse strains used in this study were of C57BL/6 background unless otherwise indicated.

The wild type strain of *M. tuberculosis* (Mtb) Erdman was used in these studies. Bacteria were cultured in 7H9 medium containing 0.05% Tween 80 and OADC enrichment (Becton Dickinson). For infections, mycobacteria were suspended in phosphate-buffered saline (PBS)-Tween 80 (0.05%); clumps were dissociated by sonication, and ∼100 CFU were delivered via the respiratory route using an aerosol generation device (Glas-Col, Terre Haute, IN). At indicated time points mice were treated with 200mg/kg of Linezolid (LZD). These drugs were prepared in 0.5% carboxymethyl cellulose (CMC) and Polyethylene glycol 300 solution as the vehicle. Cohorts of mice were treated with anti-IL-1R1 antibody (InVivoMab, anti-mouse IL-1R, Clone JAMA147, BioXcell), either alone or in combination with LZD. 0.5% CMC and Polyethylene glycol was used as vehicle control. All antibiotic treatment was done by daily oral gavage. Anti-IL-1R1 (100ug/mouse/0.2mL) was administered every alternate day by subcutaneous and intraperitoneal route.

### Macaque pharmacokinetic study and analytical method

Uninfected macaques designated for other studies (n=3) were given 20mg/kg or 40mg/kg by oral gavage and plasma acquired at 0, 5, 19, 15, 20, 25 and 30 hours post LZD administration. High pressure liquid chromatography coupled to tandem mass spectrometry (LC/MS-MS) analysis was performed on a Sciex Applied Biosystems Qtrap 4000 triple-quadrupole mass spectrometer coupled to an Agilent 1260 HPLC system to quantify LZD in macaque plasma. LZD chromatography was performed on an Agilent Zorbax SB-C8 column (2.1×30 mm; particle size, 3.5 µm) using a reverse phase gradient elution. Milli-Q deionized water with 0.1% formic acid was used for the aqueous mobile phase and 0.1% formic acid in acetonitrile for the organic mobile phase. Multiple-reaction monitoring (MRM) of parent/daughter transitions in electrospray positive-ionization mode was used to quantify the analytes. Sample analysis was accepted if the concentrations of the quality control samples were within 20% of the nominal concentration. Data processing was performed using Analyst software (version 1.6.2; Applied Biosystems Sciex).

Neat 1 mg/mL DMSO stocks for all compounds were serial diluted in 50/50 Acetonitrile water to create standard curves and quality control spiking solutions. 20 µL of neat spiking solutions were added to 20 µL of drug free plasma or control tissue homogenate, and extraction was performed by adding 180 µL of Acetonitrile/Methanol 50/50 protein precipitation solvent containing the internal standard (10 ng/mL verapamil & deuterated LZD-d3). Extracts were vortexed for 5 minutes and centrifuged at 4000 RPM for 5 minutes. 100 µL or supernatant was transferred for LC-MS/MS analysis and diluted with 100 µL of Milli-Q deionized water. Rhesus macaque plasma (Lithium Heparin, Bioreclamation IVT, NY) was used as a surrogate to cynomolgus macaque plasma to build standard curves. LZD-d3 internal standard and verapamil were purchased from Toronto Research Chemical. The lower and upper limits of quantitation (LLOQ and ULOQ) were 1 ng/mL and 50,000 ng/mL respectively. The following MRM transitions were used for LZD (338.00/235.00), LZD-d3(341.20/297.20), and verapamil (455.40/165.20).

### Macaques, infection and treatment

All housing, care, and experimental procedures were approved by the University of Pittsburgh School of Medicine Institutional Animal Care and Use Committee (IACUC). Examination of animals was performed in quarantine to assess physical health and confirmation of no previous *M. tuberculosis* infections as previously described (23). Cynomolgus macaques (*Macaca fascicularis*) (N=10) were purchased for this study from (Valley Biosystems). Bone marrow control (non-drug treated) samples were taken from Mtb-infected cynomolgus macaques in unrelated ongoing studies (N=5). For the current study, all 10 animals were infected bronchoscopically with 12 CFU of Mtb strain Erdman. After active disease developed (3-5 months), NHPs were randomized to LZD only (N=5) or LZD+IL-1Rn (N=5) treatment groups. For randomization, macaques were paired based on total FDG activity in lungs (a surrogate for total thoracic CFU), and then assigned to treatment by coin flip (20). LZD was administered twice a day orally with food (30mg/kg), while IL1-Rn was given at 2mg/kg once each day by subcutaneous injection. All animal data are provided in Supplementary Table 1. Medication compliancy was monitored at every administration and pharmacokinetic analysis performed at select times. Drug treatments were administered for 4 weeks prior to necropsy. Bronchoalveolar lavages (BAL) were performed prior to drug-treatment and 3 weeks-post start of treatment for CFU, flow cytometry and multiplex assays.

### Macaque PET/CT Imaging

Positron emission tomography (PET) with computed tomography (CT) imaging was performed with 2-deoxy-2-[^18^F]-D-deoxyglucose (FDG) throughout the study as previously described (19). Serial scans were performed throughout the study to track disease progression and changes during drug treatment. Total FDG activity of the lungs was measured over the course of infection and drug treatment as previously described (19). Granulomas identified on scans were denoted, measured (mm) and standard uptake values (SUVR) were determined to assess metabolic activity, a readout for inflammation. SUVR values were normalized to muscle and SUVR and size measurements were determined at each scan over time to compare pre-and post-drug treatment. Each animal was scanned prior to necropsy to identify granulomas for matching at necropsy; granulomas ≥ 1mm are distinguishable by PET/CT.

### Macaque Necropsy

Necropsies were performed as previously described. In short, multiple tissues (granulomas, lung lobes, thoracic lymph nodes, peripheral lymph nodes, liver, spleen, bone marrow) were excised and homogenized into single-cell suspensions for assessment of bacterial burden and immunological assays. Granulomas were individually excised (PET/CT identified and others not identified on scans) and split (size permitting) with one-half for homogenization and single cell suspension and the other half processed for histological analysis. Bone marrow samples were obtained from the sternum, with a portion sent for histological analysis while single cell suspensions were acquired as previously described (24). Bacterial burden was assessed from each tissue by plating serial dilutions on 7H11 agar plates and incubated at 37°C in 5% CO_2_ for 21 days before enumeration of Mtb CFU.

### Flow cytometry and Immunoassays

#### Mice

Single cell suspensions were prepared from the infected mouse organs. Briefly, lung tissue was digested with Collagenase type IV/DNaseI and passed through 40 μm cell strainers to obtain single cell suspension. Red blood cells were lysed using Tris-buffered Ammonium Chloride (ACT) Non-specific antibody binding sites were blocked by Fc-Block CD16/32 (Clone 93, cat. no. 101319) and the cells were stained with anti-CD3-PE (Clone 17A2, cat. no. 100205), anti-CD11b-PerCP Cy5.5 (Clone M1/70, cat. no.101227), anti-Ly-6G-FITC (Clone 1A8, cat. no.127605), anti-Ly-6C-PE (Clone HK1.4, cat. no.128007), anti-Gr1-APC (Clone RB6-8C5, cat. no.108411), anti-Ter119-PE (Clone TER119, cat. no.116208), anti-CD71-FITC (clone RI7217, cat. no. 113806). Antibodies were purchased from BioLegend. All analyses were performed on live cells only after staining them with fixable live dead stain conjugated with eFlour780, purchased from eBiosciences. All the staining was done according to the manufacturer’s instructions. Lung, spleen and bone marrow cells were surface stained for 30 minutes at room temperature, fixed for 20 minutes at 4^0^C using the Cytofix buffer (BD-Biosciences, cat. no. 554655). Data were acquired in a BD LSRII flow cytometer in the flow cytometry core facility at UMASS medical school and analyzed with FlowJo Software (Treestar, Inc.). Gating strategies are provided in applicable figures.

#### Macaques

Single cell suspensions acquired from homogenization of granulomas, lung lobes and lymph nodes were subjected to intracellular cytokine staining (ICS). Prior to staining, cells were incubated in RPMI 1640 containing 1% HEPES, 1% L-glutamine, 10% human AB serum, and 0.1% brefeldin A (Golgiplug; BD Biosciences) for 3 h at 37°C in 5% CO_2._ After viability staining (Invitrogen), surface antigens and intracellular cytokines were assessed using standard protocols. Surface markers include CD3 (SP34-2; BD Pharmingen), CD4 (L200; BD Horizon), CD8 (RPA-T8; BD Horizon) for T cells and CD11b (ICRF44; BD Pharmingen), CD206 (19.2; BE Pharmingen) for macrophages/neutrophils. Calprotectin (27E10; ThermoFisher) was stained intracellularly to identify neutrophils. For bone marrow, single cell suspensions underwent red blood cell lysis (BD Pharm Lyse) before incubation in alpha-MEM + 10% Stasis^TM^ FBS (Gemini Bio-Products) + MitoTracker^TM^ Red CMXRos (Invitrogen) for 30 minutes to stain for membrane potential of mitochondria. Cells were then stained for viability (Invitrogen) and surface stained to distinguish erythroid progenitor populations by CD34 (581; Biolegend), CD235a (HIR2; BD Pharmingen), CD71 (L01.1; BD Pharmingen) and CD45 (D058-1283; BD Pharmingen). All samples were acquired on an LSR II (BD) and analyzed with FlowJo Software (Treestar, Inc.). Gating strategies are provided in applicable figures.

For the multiplex assays, all samples and supernatants were stored at −80°C from time of necropsy until time of assay. Five representative granuloma supernatants were randomly selected from each animal using JMP Pro v12 (SAS Institute Inc.). Supernatants were thawed and filtered with a 0.22uM syringe filter to remove infectious bacteria and debris, then kept on ice throughout the assay. For BAL samples, supernatants were concentrated using regenerated cellulose centrifugal filter tubes (3,000 NMWL, Millipore Sigma) to a final 10X concentration (5mL to 0.5mL). Both granuloma and BAL supernatants were evaluated with a ProcartaPlex multiplex immunoassay (Invitrogen) that assesses thirty cytokines and chemokines specific for NHPs. We followed the manufacturer’s protocol with one modification in which we diluted the standard out an extra dilution to increase the range of detection. Results were analyzed by a BioPlex reader (BioRad).

### Histopathology and immunofluorescence

#### Mice

Lung tissues were fixed in 10% buffered formalin and embedded in paraffin. Five micrometer-thick sections were stained with hematoxylin and eosin (H&E). Tissue staining was done by the Diabetes and Endocrinology Research Center histopathology core facility at the University of Massachusetts Medical School or immunology core facility of Albany medical college, NY. Brightfield images were acquired in Abaxis VETSCAN HD microscope.

Paraffin embedded lung tissue sections were cut at 5 μm thickness, mounted on ultraclean glass slides covered in silane, deparaffinized, then dehydrated and rehydrated using the following steps: Ethanol solutions (30, 50, 70, 90, 95 and 100 % for 3 min each), xylenes (2 different solutions for 10 min each) and ethanol solutions (100, 95, 90, 70, 50 and 30 for 3 min each). The slides were washed once in Tris buffer saline (TBS) for 5 min. Slices were subjected to antigen retrieval by boiling in sodium citrate buffer at pH=6.0 for 20 min and incubated in 0.1% Triton-X 100 for 5 min. Slices were removed and allowed to equilibrate to room temperature for at least 20 min and rinsed with distilled water. Tissue sections were blocked (blocking solution; 0.5 M EDTA, 1% BSA, in PBS) and incubated overnight in primary antibodies against the proteins related to our studies. Sections were stained for nuclei (DAPI, blue staining), anti-mouse CD3e (cat.no. ab16669, green staining), anti-mouse Ly-6G (clone 1A8, cat. no. 127602) to identify neutrophil granulocytes (Cy3, red staining). As controls, pre-immune serum and isotype matched controls were used. After incubation, the tissues were washed several times with sterile TBS at room temperature and incubated in the respective secondary antibodies (anti-rabbit conjugated to Alexa-488, anti-rat conjugated to Cy3) for at least 2h at room temperature. Tissue sections were mounted using Prolong Gold Antifade reagent (Invitrogen, grand Island, NY) with DAPI, and the tissue sections were examined in ECHO Revolve 4 microscope. Isotype matched control antibodies were used for checking antibody specificity.

#### Macaques

As previously described, samples acquired at necropsy were formalin fixed, paraffin embedded, cut into ∼5μm serial sections and stained with H&E (23). A study-blinded veterinary pathologist assessed and characterized each granuloma, indicating size, type, distribution and cellular composition. Sterna for bone marrow analysis were excised and formalin fixed then transected longitudinally into 1 to 2 sternebral unites and placed in Cal-Ex Hydrochloric acid decalcification solution for 2-4 hours. Upon removal, specimens were washed and trimmed to test for adequate mineral removal, then submitted for routine tissue processing with other tissue specimens as described above.

### Statistical Analysis

#### Mouse studies

Equal variance between samples was assessed by Brown-Forsythe test. Experiments in which variances were equivalent were analyzed by one-way ANOVA with Sidak’s multiple comparisons test. Those with unequal variances were analyzed by Welch ANOVA and Dunnett’s multiple comparisons test. P values < 0.05 was considered significant.

#### Macaque studies

Nonparametric U tests (Mann-Whitney) were performed for two-group comparisons and Kruskal-Wallis tests were performed for three-group comparisons as indicated on data sets with non-normal distributions. Wilcoxon signed rank tests were performed for matched pairs. Fisher’s exact test was run on any categorical data. P values < 0.05 were considered significant. Total lung FDG activity was log_10_-transformed. Note that for any log-transformed data, a log_10_(x + 1) transform was implemented so that zeroes were not excluded from the graphs or analyses. A two-way ANOVA was utilized to test whether treatment or time (or an interaction between these factors) had an effect on total lung inflammation. Time was found to be a statistically significant effect (p=0.0046); therefore, Dunnett’s multiple comparison tests were then used to compare each time point to pre-drug treatment within each treatment group. For bivariate data, a linear regression was used to test the linear relationship between the variables and the R^2^ and p-value for the F-test are reported in the accompanying figure legend. Statistical analyses were performed in GraphPad Prism 8 (GraphPad Software, San Diego, CA). A regression equation created from control animals from previous studies was used to create a 95% prediction interval for total thoracic bacterial burden using total lung FDG activity on the scan just before drug treatment (20). The lower and upper bounds and the mean of this prediction interval were subtracted from each animal’s total CFU to estimate change in CFU over the course drug treatment. All statistical analyses are referenced in the corresponding figure legends.

## Results

### IL-1 receptor blockade reduces inflammation in mouse models of TB disease

Given the complex role played by IL-1 during TB, we initially sought to determine the effect of inhibiting this cytokine during TB disease in mice. We used a blocking antibody to the murine IL-1 receptor type 1 (αIL-1R1) since this reagent has been shown to alter TB disease in mice (25), and we previously found that recombinant human IL-1Rn (Anakinra) has little effect in this mouse model. Two mouse strains were employed to assess the effects of these treatments in animals with different amounts of IL-1 activity. In relatively resistant C57BL/6 animals, mature IL-1 production is controlled by IFNγ-dependent nitric oxide production, whereas unregulated IL-1 drives inflammatory disease characterized by increased number of neutrophils (PMNs), bacterial burden (CFU) in the lungs and significant weight loss observed in Mtb-infected *Nos2*^-/-^ mice that lack this regulatory pathway (9, 10).

αIL-1R1 was administered to the animals between days 14 and 28 post infection after the onset of adaptive immunity. In both resistant (C57BL/6) and susceptible (Nos2^-/-^) mice, αIL-1R1 treatment significantly reduced PMN numbers in the lungs. In addition, this treatment reversed the weight loss observed in *Nos2*^-/-^ animals (Fig. 1A-B). Somewhat surprisingly, this regimen also modestly reduced the mean bacterial burden in lungs and spleens, with this reduction meeting statistical significance in the spleens of *Nos2^-/-^* mice (Fig. 1C-D). This generally beneficial effect of αIL-1R1 treatment was consistent with qualitatively improved histopathological disease (Fig. 1E). By no metric did this αIL-1R1 treatment exacerbate disease in either mouse strain.

**Figure 1.**
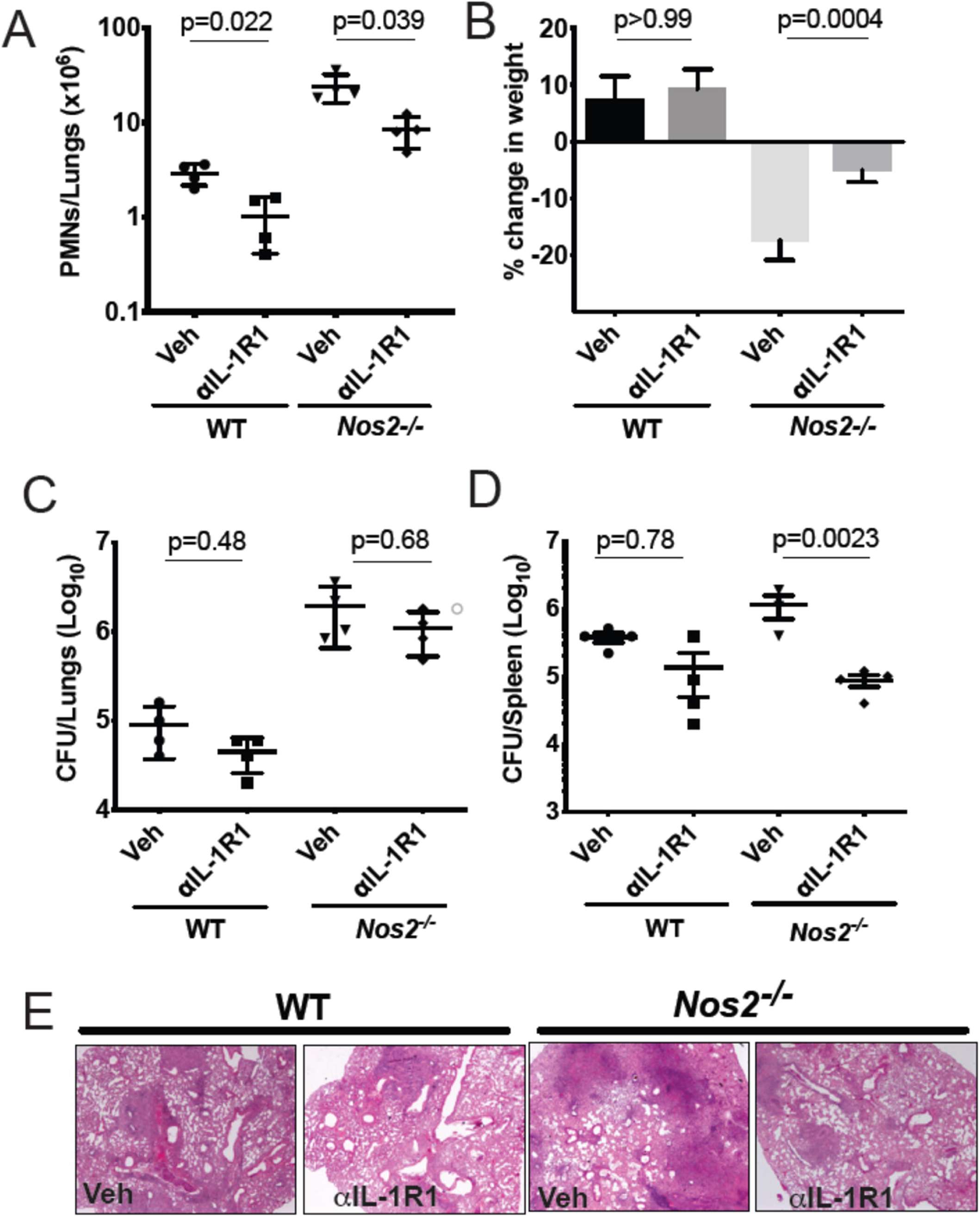
IL-1 inhibition in susceptible mice reduces inflammation with no effect on lung bacterial burden. Wild type C57BL/6 and *Nos2^-/-^* mice were infected with Mtb Erdman for 2 weeks and treated with anti-IL-1R1 (αIL-1R1) for the subsequent 2 weeks. (A) Lung neutrophil (PMN) infiltration was quantified by flow cytometry; (B) Weight loss and (C) CFU in lung and (D) spleen are shown. Data shown (mean ± SD) are representative of two independent experiments. One-way ANOVA with Sidak’s multiple comparisons-test was used. N=4 mice per treatment cohorts. (E) Histopathology analysis of a single lung lobe from WT or *Nos2-/-* mice was performed by Hematoxylin-eosin (H&E) staining.

### IL-1 blockade alleviates lung inflammation and hematopoietic suppression during LZD treatment in mice

The IL-1R1 blocking antibody was next tested in combination with LZD to determine whether the efficacy or toxicity of the antibiotic was altered. C3HeB/FeJ mice, which are relatively susceptible to Mtb and develop histopathological lesions that more closely resemble human disease, were used for these studies. To model established disease, mice were treated between days 28 and 46 post-infection with vehicle alone, LZD, αIL-1R1 or a combination of the two. As previously reported, LZD was effective in this model, reducing lung neutrophil numbers, bacterial burden and weight loss (Fig. 2 A-D) (26). The addition of αIL-1R1 to this regimen further reduced lung neutrophil numbers. IL-1 blockade had very little effect on bacterial burdens in lung or spleen, whether given alone or in conjunction with LZD. The mean CFU burden in αIL-1R1-treated animals was within 2.3-fold of the untreated groups, and only the αIL-1R1-associated decrease in the spleens of LZD treated animals approached significance. As IL-1β production in response to both Mtb infection (9) and LZD treatment (16) depends largely on the NLRP3 inflammasome, we also investigated a regimen in which LZD and a small molecule NLRP3 inhibitor (MCC950) was administered between days 56 and 77 post-infection. As observed with αIL-1R1, the addition of MCC950 reduced PMN numbers in the lung, relative to LZD alone, and did not significantly alter bacterial killing (Fig. 2 E-G). Using a more rapid treatment protocol and C57BL/6 mice with genetic deficiencies in Caspase 1 or NLRP3, we confirmed that the antimicrobial activity of LZD was unaffected by inflammasome activation (Supplementary Fig. 1A-D). Mice were treated between days 14 and 28 post-infection and the effect of LZD on bacterial burden and lung neutrophil number was at least as large in the knockout animals as the wild type controls.

**Figure 2.**
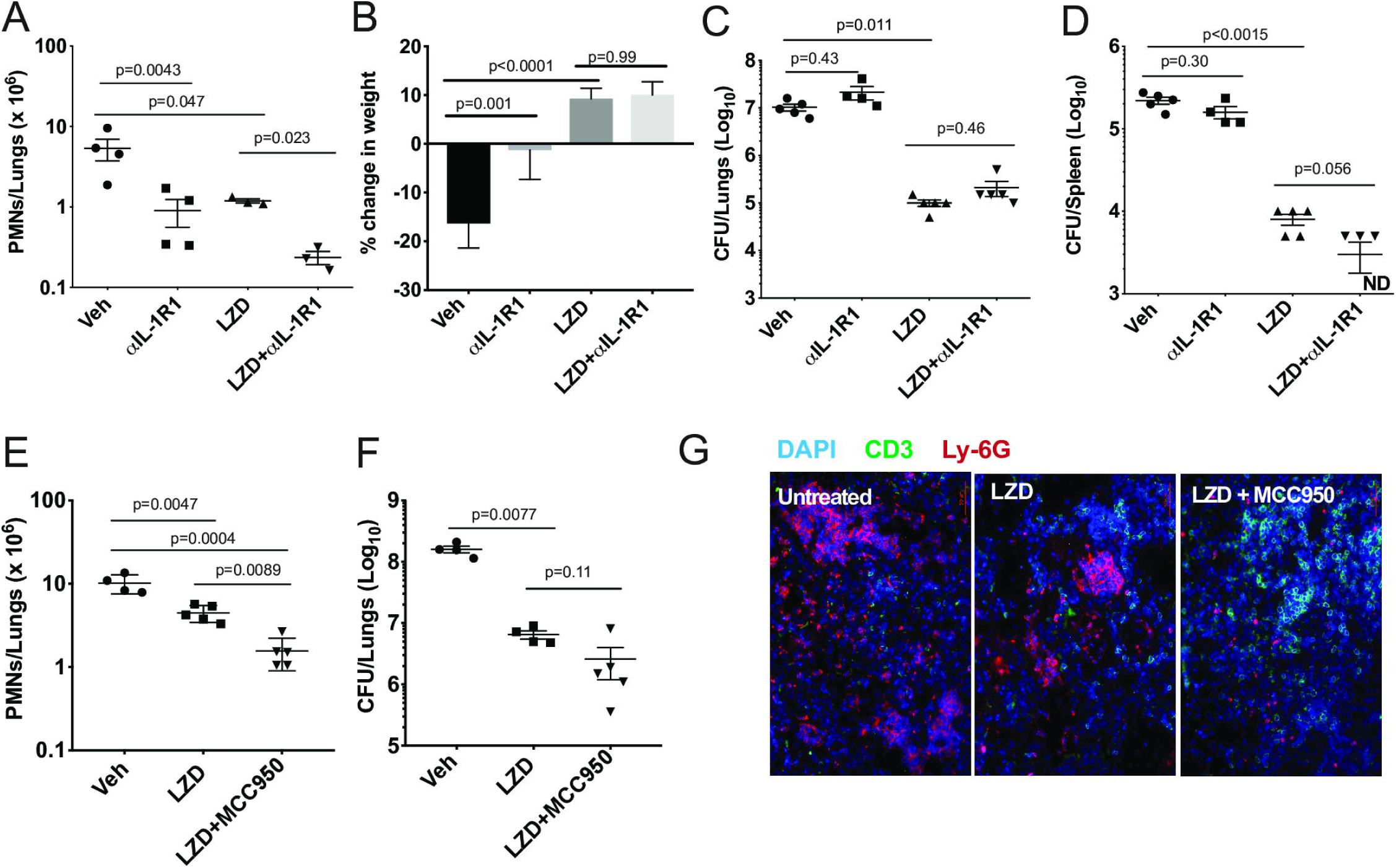
IL-1R1 blockade combined with Linezolid ameliorates TB disease. (A-D) C3HeB/FeJ mice were infected with Mtb Erdman for 4 weeks and treated with linezolid (LZD) and/or anti-IL-1R1 (αIL-1R1) for the following 18 days. (A) Lung neutrophils were quantified by flow cytometry; (B) Percent change in body weight is shown and (C) Bacterial burden in the lung and (D) spleen were quantified as CFU. Data shown (Mean ± SD) are representative of two independent experiments. Welch ANOVA with Dunnett’s post-test was used to calculate statistical significance where each treatment group was compared to the vehicle as control group. N=3-5 mice per treatment group. (E-G) C3HeB/FeJ mice were infected with *M. tb* Erdman for 8 weeks and treated with linezolid (LZD) and/or an inhibitor of the NLRP3 inflammasome (MCC950) for the following 21 days. (E) Lung neutrophils were quantified by flow cytometry. (F) Bacterial burden in the lung was quantified by CFU. Data shown (Mean ± SD) are from one experiment. One-way ANOVA with Tukey’s multiple comparison test was used to calculate the p-value. N=4-5 mice/group. (G) Representative immunofluorescence images of the lungs from different treatment groups. Cell nuclei stained with DAPI (blue), T-cells stained with anti-mouse CD3ε (green) and neutrophils stained with anti-mouse Ly-6G (red).

Next, we evaluated the hematopoietic suppression caused due to LZD treatment in the bone marrow and spleens of C3HeB/FeJ mice. In small animals, bone marrow serves as the primary site of hematopoiesis, but during infection or stress, the spleen functions as an extramedullary hematopoietic organ to compensate for the increasing demand of blood cells in the periphery(27). As LZD did not alter the myeloid cells in the bone marrow (data not shown), and previous reports have shown its effects on erythropoiesis (16), we investigated the effect of this oxazolidinone on erythroid lineage cells in both these organs. Using flow cytometry, erythroid progenitors can be divided into a progression of precursors: pro-erythrocyte (ProE), EryA, EryB, and EryC (Fig. 3A). These populations were quantified in each tissue of Mtb-infected animals treated with LZD and/or αIL-1R1. In both bone marrow and spleen, LZD had a profound effect, nearly eliminating early erythroid progenitors of the ProE, EryA and EryB classes (Fig. 3B-E).

**Figure 3.**
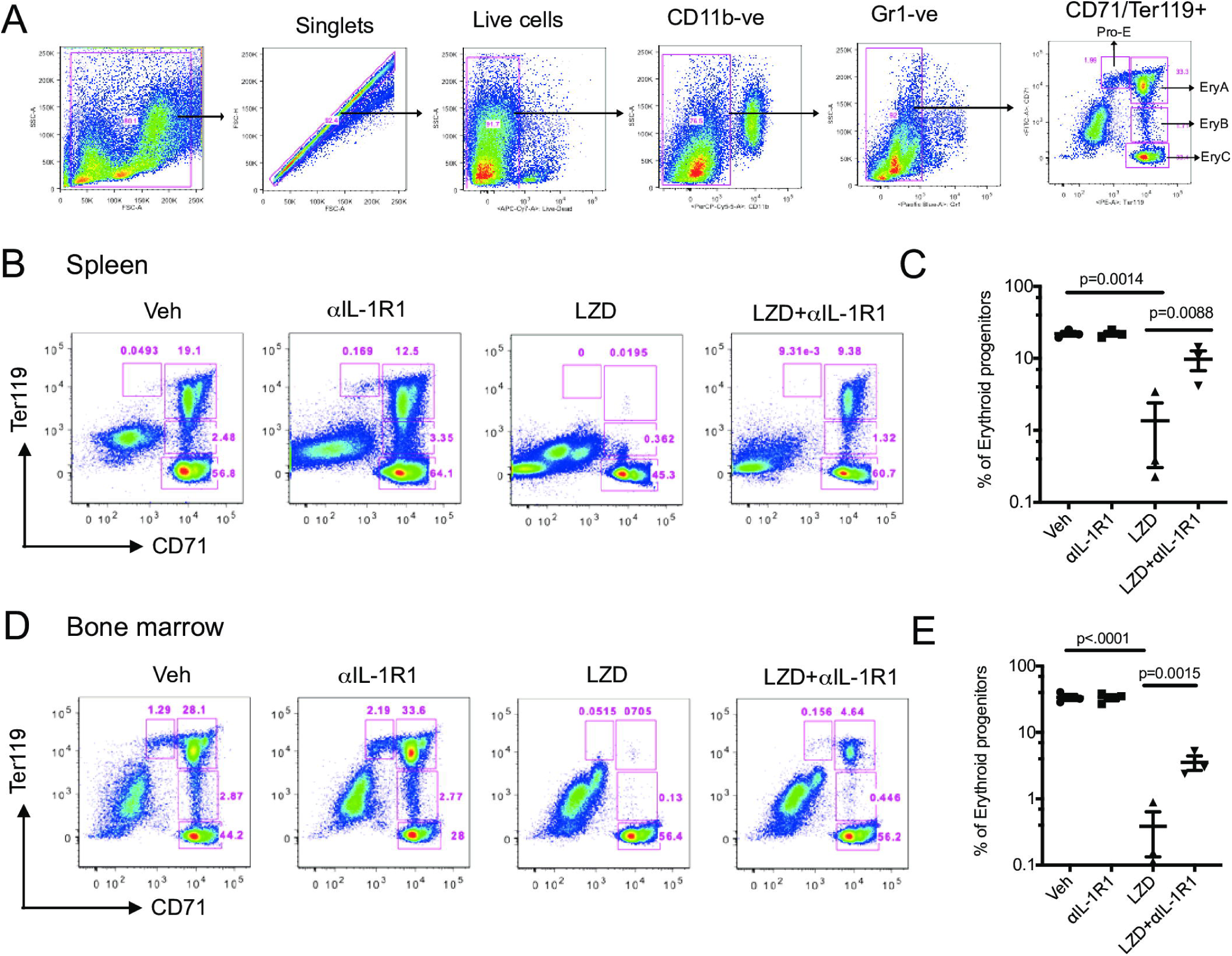
Erythropoiesis associated with LZD treatment was partially relieved after IL-1R1 blockade. C3HeB/FeJ mice were infected with Mtb Erdman for 4 weeks, and treated with LZD either alone or in combination with anti-IL-1R1 antibody for the subsequent 18 days. (A) Gating strategy for detecting erythroid progenitors is shown. Erythroid progenitors in the spleen (B-C) and bone marrow (D-E) were quantified by flow cytometry as described in Materials and Methods. Representative flow cytometry plots and percent changes in the early erythroid progenitors (Pro-Ery, EryA, EryB) among different treatment groups are shown. Data shown (Mean ± SD) are from two independent experiments. One-way ANOVA with Dunnett’s post-test was applied to calculate the p-value by comparing the mean of each group with that of LZD treated group. N=3-5 mice per group.

Simultaneous treatment with αIL-1R1 largely reversed the effect of LZD in the spleen, restoring these immature precursors to approximately half of their untreated levels. While αIL-1R1 also significantly increased the number of erythroid precursors in the BM, the suppression of LZD toxicity was less pronounced at this site.

In sum, studies in the mouse model indicated that the addition of αIL-1R1 to an LZD regimen could reduce the number of lungs PMN and ameliorate hematopoietic toxicity, while not compromising antimicrobial activity. These observations justified further studies in a non-human primate model.

### Changes in TB disease in macaques treated with LZD and HDT by PET/CT

Previously we published the efficacy and pharmacokinetics of LZD in cynomolgus macaques treated for 8 weeks with a single daily dose of 30 mg/kg (14). To adhere to FDA guidelines for LZD administration at the time we initiated this study and reproduce clinical exposure at 600 mg twice daily (b.i.d.), we shortened treatment duration to 4 weeks and increased the dosing frequency to 30 mg/kg b.i.d. Dose finding studies were carried out in uninfected cynomolgus macaques to ensure that adequate drug concentrations similar to those achieved in patients at 600 mg b.i.d. were reached in the blood (Supplementary Fig. 2).

To assess this LZD regimen and HDT with IL-1Rn, we infected 10 cynomolgus macaques with Mtb strain Erdman and monitored development of active TB disease (Supplementary Table 1 and Supplementary Fig. 2). Infected macaques were then randomized to a 4-week drug regimen of LZD (n=5) or LZD+IL-1Rn (n=5) (Supplementary Fig. 2). Our model has the advantage of tracking disease progression throughout infection and treatment using serial ^18^F-FDG PET CT scans (14, 28). Lung inflammation quantified by total FDG activity in the lungs is correlated with total thoracic bacterial burden as previously described (19). Here we quantified total lung FDG activity before and during treatment. 3D rendered images provide a visual for changes in overall lung inflammation in the “best” and “worst” animals for both treatment groups (Fig. 4A). Quantification of total lung FDG showed no significant difference in inflammation at time of necropsy between LZD and LZD+IL-1Rn treatment groups (Fig. 4B), however a slight change in slope with IL-1Rn treatment was observed. To expand on this finding and test if LZD+IL-1Rn reduced total lung FDG activity more effectively than LZD alone over time, we compared the change in FDG activity between 0, 2 and 4 weeks post treatment. In LZD only treated macaques, there was no significant decrease in total lung FDG activity after 2 (p=0.0857) and 4 weeks of treatment (p=0.2876) (Fig. 4B). In contrast, LZD+IL-1Rn treatment resulted in a significant reduction in total lung FDG activity after 2 and 4 weeks (p=0.0242, p=0.0237 respectively). To assess whether these changes in lung inflammation were reflected in individual granulomas, we used PET CT scans acquired pre-treatment and at 4 weeks post-treatment to determine changes in granuloma physical size (mm) and metabolic activity as Standard Uptake Value (SUVR) of FDG per granuloma (Fig. 4C-D) (19). SUVR indicates the inflammatory state of each individual granuloma, allowing us to track specific lesions throughout treatment for changes in activity. After treatment, while the majority of granulomas decreased in size and SUVR, there was variability within an animal and between animals and no significant differences between LZD and LZD+IL1-Rn treated animals. These data indicate that blocking of IL-1R in combination with LZD reduces total lung inflammation more efficiently than LZD alone.

**Figure 4.**
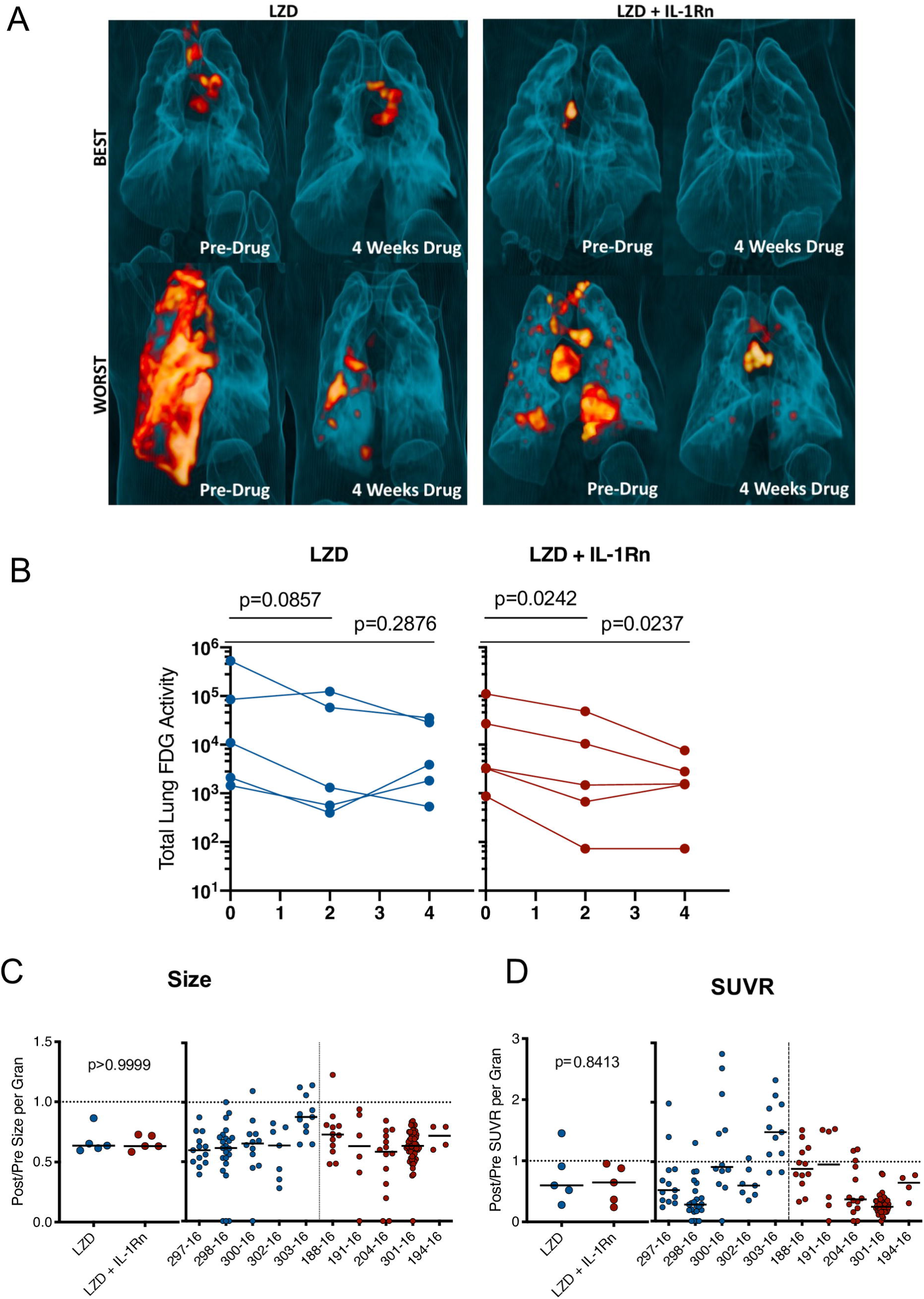
HDT reduces granuloma inflammation by PET/CT in macaques. Cynomolgus macaques were infected with *Mtb* Erdman for approximately 4 months (see Supplementary Table 1) and randomized to treatment with LZD or LZD+IL-1Rn for an additional 4 weeks. PET CT scans were performed pre-treatment 0, 2 and 4 weeks post-treatment with 4 week scans as the last prior to necropsy. (A) 3-D renderings of PET/CT scans from pre-treatment and 4 weeks post-treatment are depicted, with “best” and “worst” of each group referring to TB disease prior to drug administration. (B) Total lung FDG activity of each macaque throughout treatment with LZD (left) or LZD+IL-1Rn (right). Two-way ANOVA with Dunnett’s adjusted p-values are reported. (C) Individual granulomas were identified pre-treatment and tracked post-treatment by PET/CT. Change in size (by CT) was determined for granulomas from each animal; the median (left) change in granuloma size from each animal and individual granulomas per animal (right). (D) Standard uptake value (SUVR) of ^18^F-FDG was calculated for each granuloma, representing inflammation. The median change in SUVR of all granulomas (left) and change in SUVR of individual granulomas from each animal (right) are shown. Mann-Whitney tests determined p values for (C) and (D), with p<0.05 considered significant.

### Addition of IL-1Rn does not alter LZD bacterial clearance

IL-1Rn does not have direct bactericidal activity, therefore addition of IL-1Rn to LZD treatment should not directly influence bacterial killing and burden. However, to ensure immunomodulation with IL-1Rn did not have detrimental effects on host antibacterial immune responses, we performed comprehensive bacterial burden analysis at necropsy as previously described (20). In short, we acquired individual granulomas identified by PET CT as well as other TB pathologies, thoracic lymph nodes, uninvolved lung tissue and extrapulmonary lesions for bacterial plating and CFU determination. Total thoracic CFU (lung + lymph nodes) was not significantly different between LZD and LZD + IL-1Rn in macaques (p = 0.1508). Similarly, separate analysis of lung CFU excluding thoracic lymph nodes did not demonstrate significant differences between groups (p=0.5476) (Fig. 5A). We did not have untreated control macaques available in this study, therefore we provide total thoracic CFU for 3 historical control animals that were similarly infected and necropsied at a similar time point; these data serve as reference for expected CFU at this time point and were not included in statistical analyses. While the treatment groups had similar frequencies of sterilized granulomas (LZD = 68.42%, LZD+IL-1Rn = 72.22%), it is important to note that the majority of granulomas were sterilized during the short four-week course of LZD or LZD+IL-1Rn treatment. We previously reported that a 2-month lower dose regimen of LZD could reduce bacterial burden compared to untreated controls, with ∼80% sterilized granulomas in treated animals and ∼20% sterilized in untreated (14). This supports that high dose LZD as a single drug is effective at killing bacteria even in a short (4 week) regimen. The lymph nodes are a recently identified reservoir for Mtb and a hub for immune cell interactions (29), therefore we assessed HDT effects on bacterial burden in thoracic lymph nodes after treatment. We saw no significant difference in overall lymph node CFU nor CFU per lymph node (Fig. 5C), indicating addition of IL-1Rn did not measurably alter LZD efficacy or antibacterial immunity in lymph nodes. To better understand the efficacy of high dose LZD for 4 weeks, we estimated the pre-treatment total thoracic CFU for each macaque using the total lung FDG by PET CT as previously described (20), and compared the true bacterial burden post-treatment against the estimated pre-treatment value. This analysis allows us to estimate the magnitude of LZD bacterial killing during LZD and LZD+IL-1Rn treatment. All macaques had lower total thoracic CFU at necropsy compared to the estimated total thoracic CFU prior to treatment start and on average the total CFU was ∼100-fold lower than estimated CFU prior to treatment (Fig. 5D). Our data indicate that while IL-1Rn did not significantly enhance bacterial killing, it did not impair LZD-mediated bacterial clearance, an important observation for HDT development.

**Figure 5.**
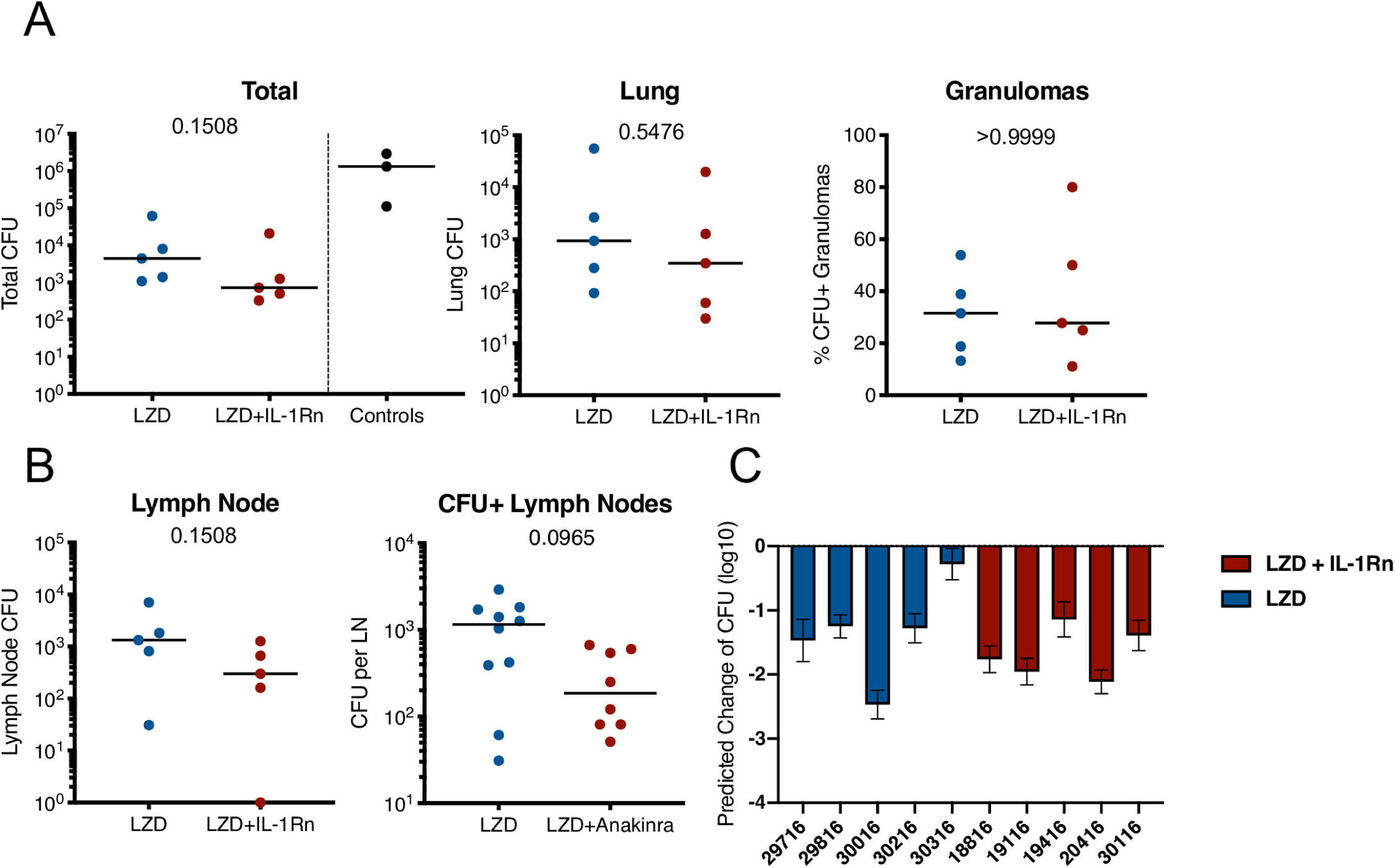
Addition of IL-1Rn does not modify efficacy of LZD-bacterial killing. Bacterial burden is shown after 4 weeks of LZD (blue) or LZD+IL-1Rn (red) treatment. (A) Total thoracic (lung + lymph nodes) and lung CFU; each data point represents one macaque. Total thoracic CFU from 3 similarly infected untreated historical control macaques (black) are included as reference for untreated CFU at this time point, but excluded from statistical analysis. (B) The percent of all lung granulomas that were CFU+ per monkey. (C) CFU of all thoracic lymph nodes with data points representing CFU from one macaque. CFU+ lymph nodes show the CFU from each thoracic lymph node in any animal that was Mtb+. For (A) and (B), p values were determined by Mann-Whitney test. (C) PET/CT scans from pre-treatment were used to model predicted CFU prior to HDT initiation and compared to actual CFU determined at time of necropsy. The predicted change in CFU was determined by: Actual total thoracic CFU post-treatment– Predicted total thoracic CFU interval (predicted from pre-drug-treatment scan). Bars represent mean of prediction intervals and error whiskers represent the lower and upper bounds of the difference between final total CFU and predicted CFU prior to drug treatment.

### Lack of LZD-induced bone marrow suppression in macaques

In humans, LZD is associated with host toxicities during extended treatment periods of greater than 4 weeks (30) resulting in the FDA limitation of LZD to less than 4 weeks for most indications. We designed our study in macaques to follow FDA-guidelines for transition to possible human trials, therefore the potential for bone marrow toxicity to occur within this timeframe in macaques was unknown. To determine whether bone marrow suppression occurred during the 4-week high dose LZD therapy and whether IL-1Rn could modulate any observable host toxicities, we isolated bone marrow at necropsy from the sternum of each macaque and assessed erythropoietic progenitor populations and mitochondrial function by flow cytometry (Fig. 6A). During homeostatic erythropoiesis, a 1:8 ratio is maintained between ProE and EryC progenitor populations. Changes in this ratio can be used to indicate disruption of erythropoiesis. In macaques, there was no significant difference in ProE:EryC ratios between the treatment groups (Fig. 6B). We also analyzed control bone marrow from untreated Mtb-infected macaques involved in other on-going studies (TB only), which showed no significant difference between ProE:EryC ratios compared to treatment groups. To determine whether LZD was affecting mitochondrial function, we stained bone marrow cells with MitoTracker^TM^ Red CMXRos which only stains mitochondria with active membrane potential, indicating function of those organelles (Fig. 6B). Regardless of treatment, the MitoTracker MFI remained similar, indicating that mitochondrial function is the same among groups. To confirm our observations, a pathologist evaluated bone marrow tissue sections and found no observable differences between TB only, LZD and LZD+IL-1Rn groups in terms of cellularity (myeloid:erythroid ratios) or abnormalities (Fig. 6C). We could not consistently identify progenitor populations in blood or spleen, therefore we performed complete blood counts (CBCs) during treatment to identify signs of bone marrow suppression (i.e., anemia, thrombocytopenia). There were no observable differences between LZD and LZD+IL-1Rn treatments (Supplementary Fig. 3). In conclusion, 4 weeks of LZD was not sufficient to induce observable bone marrow suppression in cynomolgus macaques and blocking of IL-1R did not alter bone marrow status during LZD treatment of TB.

**Figure 6.**
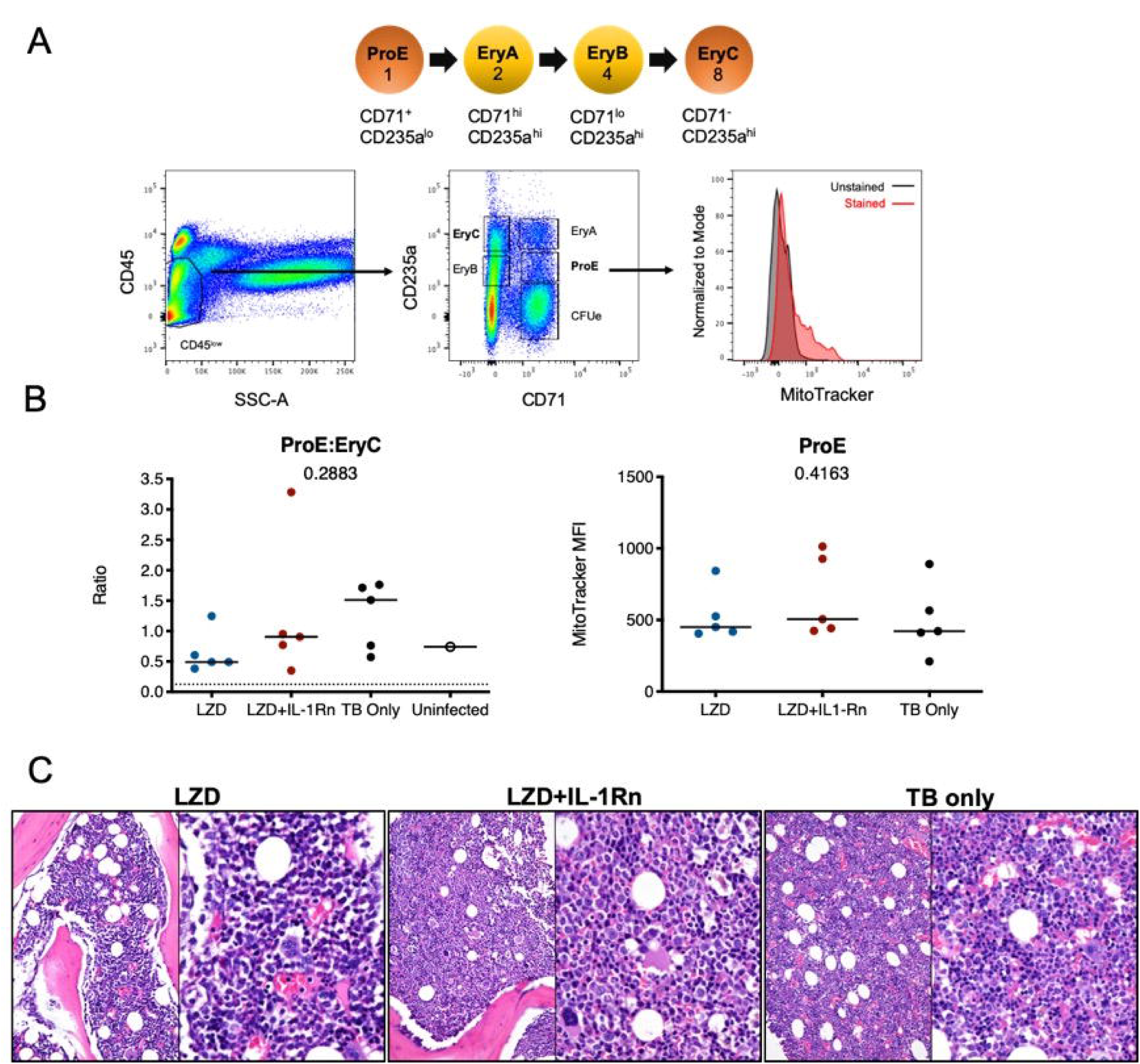
Linezolid-associated bone marrow suppression was not observed in macaques. Bone marrow was acquired from the sternum of all macaques at necropsy after 4 weeks LZD or LZD+IL-1Rn. A portion was used for flow cytometric analysis of erythroid progenitor populations and the rest for histopathology analysis. (A) A schematic of erythropoiesis indicating the stage of differentiation and corresponding expression levels of phenotyping markers (CD235a and CD71) and progenitor ratios (1:2:4:8). Included is the gating strategy to identify progenitor populations and an example of MitoTracker staining. (B) The ProE:EryC ratio (left) of each animal, including untreated TB only macaques (black) from other ongoing unpublished studies. A single uninfected animal (open circle) is shown as reference, the dotted line indicates the homeostatic ratio of 1:8. MitoTracker MFI (right) is shown for each macaque per treatment group. Kruskal-Wallis test was performed to compare the three groups (excluding uninfected macaque) and p values are shown in the graphs. (C) Representative H&E images from sternal bone marrow acquired at necropsy, 20x (left) and 40x (right) are shown.

### Reduction in neutrophil signatures in the lung during treatment

To determine whether addition of IL-1Rn to LZD treatment modulated immune populations similar to observations in the mouse model, bronchoalveolar lavages (BAL) were acquired prior to the start of treatment and 3 weeks post treatment. Innate and adaptive immune cell populations were identified and frequencies were assessed by flow cytometry (Fig. 7A). Population frequencies of CD4 T cells, CD8 T cells, neutrophils (PMNs) and macrophages remain similar between LZD or LZD+IL-1Rn, indicating that IL-1Rn did not significantly alter the frequencies of immune populations in the airways. However, we observed a possible reduction in neutrophils post treatment in both groups. Therefore we pooled LZD and LZD+IL-1Rn treated animals and compared pre and post treatment immune populations. Although macrophage populations were relatively unchanged, there was a significant reduction in neutrophil frequency in the BAL after 3 weeks of treatment with LZD, regardless of IL-1Rn (p=0.0098). To determine whether the cytokine milieu in the airways changed during treatment, a subset of 3 animals per group were chosen at random for analysis by multi-plex using 10X concentrated BAL fluid (Fig. 7B). Statistical analysis was performed on combined treatment groups to only compare pre- and post-treatment changes. We assessed IL-1β and IL-1RA and saw no significant changes after treatment (IL-1β p=0.1562, IL-1RA p=0.1250). Interferon-inducible T cell alpha chemoattractant (I-TAC) is upregulated in response in interferons and IL-1 and is generally an indicator of inflammation, which appeared to be reduced post treatment, however was not statistically significant (p=0.0625). IL-8, a neutrophil chemoattractant, was significantly reduced after treatment, mirroring the reduction in neutrophil frequency observed by flow cytometry and the mouse data. These data suggest that treatment with either LZD or LZD+IL-1Rn rapidly reduces inflammatory signatures associated with TB disease in the airways.

**Figure 7.**
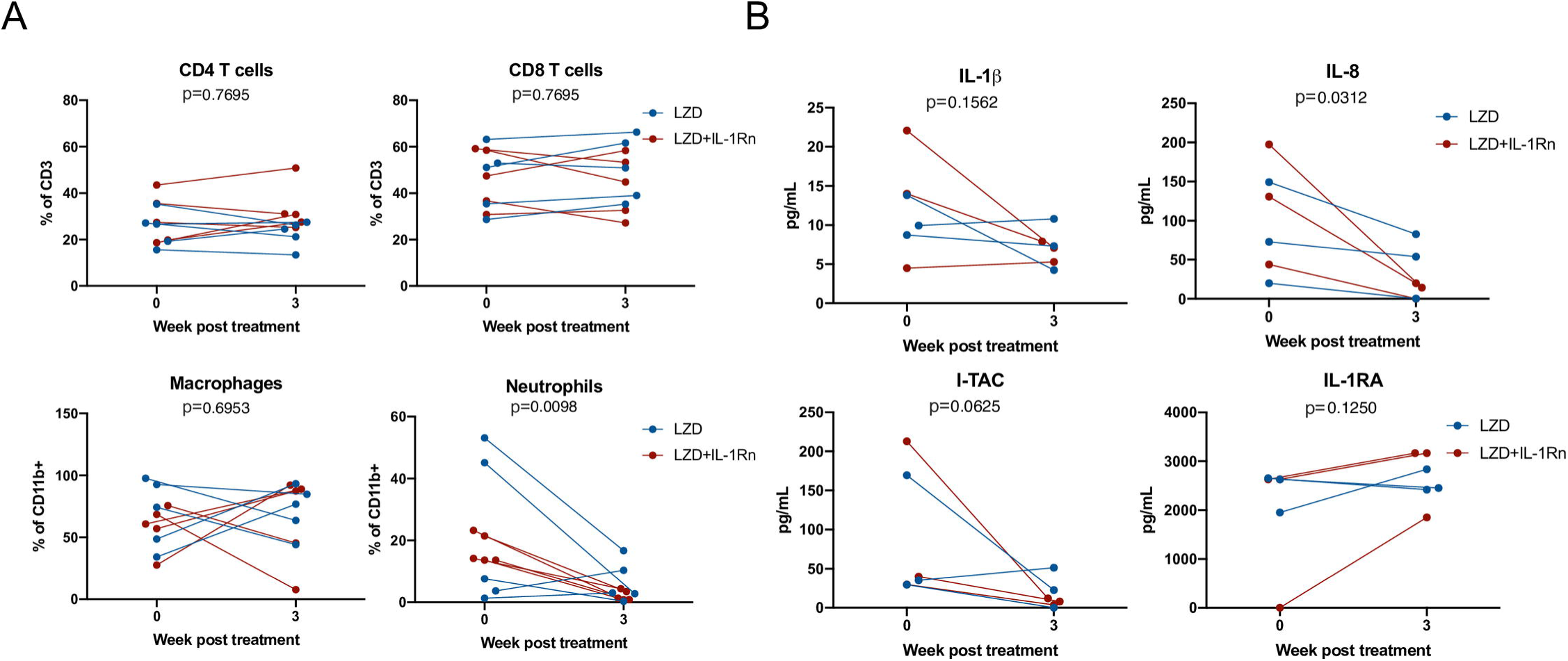
HDT reduces inflammatory signatures in BAL. Bronchoalveolar lavages (BAL) were acquired pre-treatment and 3 weeks post-treatment with LZD (blue) or LZD+IL-1Rn (red). (A) Cells were analyzed by flow cytometry to determine frequency changes in CD4 T cells (CD3+CD4+), CD8 T cells (CD3+CD8+), macrophages (CD11b+CD206+) and neutrophils (CD11b+Calprotectin+). Wilcoxon signed rank test was performed to determine differences before and after drug treatment, regardless of group (LZD and LZD+IL-1Rn combined). (B) BAL fluid was concentrated 10X and assessed by multiplex assay for changes in inflammatory cytokines and chemokines for a random subset of samples (n=3 per treatment group). Wilcoxon signed rank test was performed to determine differences before and after drug treatment, regardless of group (LZD and LZD+IL-1Rn combined).

### IL-1 blockade modulates granuloma environment and pathology

To determine whether IL-1Rn modulated immune responses at the site of infection, we chose at random 5 granulomas per animal (25 per treatment group) and performed a multi-plex analysis of granuloma supernatants (Fig. 8A). There were no significant differences in IL-1β, IL-1RA, or IL-18 levels in either treatment group. IL-2 and IL-17 are correlated with protective immune responses during TB (31) and while IL-2 levels in LZD+IL-1Rn treatment were not statistically higher (p=0.0882), a statistically significant increase in IL-17a in LZD+IL-1Rn treated animals was observed. Interestingly, G-CSF/CSF-3 levels were also significantly higher in LZD+IL-1Rn treated granulomas (p=0.0004). These results indicate IL-1Rn immune modulation occurs in the granuloma environment. While we did not see a significant difference in granuloma inflammation by SUVR between treatment groups (Fig 4C), we questioned whether the changes in cytokines could be associated with change in SUVR. Therefore, we performed linear regression analyses and found a positive correlation between IL-1β and fold change of SUVR in LZD+IL-1Rn granulomas (Fig 8B). The IL-1 pathway is associated with fibrosis, which increases in granulomas during drug treatment (32). Furthermore, IL-17 and G-CSF are also associated with fibrosis (33). To determine whether immunomodulation by IL-1Rn changed granuloma pathology, granulomas suitable for histological analysis were evaluated by a study blinded pathologist and lesions were categorized based on treatment groups and descriptive qualities (Fig. 8C). Of lesions acquired from LZD only animals, ∼71% were identified as fibrotic (n= 40/56), while those treated with LZD+IL-1Rn had ∼85% fibrotic lesions (n=39/46) with no significant difference in frequency of fibrotic lesions between treatment groups. We also compared the frequency of granulomas that were deemed necrotizing or non-necrotizing (“other” indicates neither categorization). Granulomas from LZD+IL-1Rn treated animals had significantly more non-necrotizing granulomas (∼74%, n=34/46), compared to those from LZD alone treated macaques (∼46%, n=26/56). The proportion of necrotizing granulomas was significantly lower in the LZD+IL-1Rn treatment group (13.04%, n=6/56) compared to LZD alone (30.36%, n=17/56) (p=0.0151). These data suggest that addition of IL-1Rn influences inflammation and antibiotic-associated pathology dynamics of granulomas.

**Figure 8.**
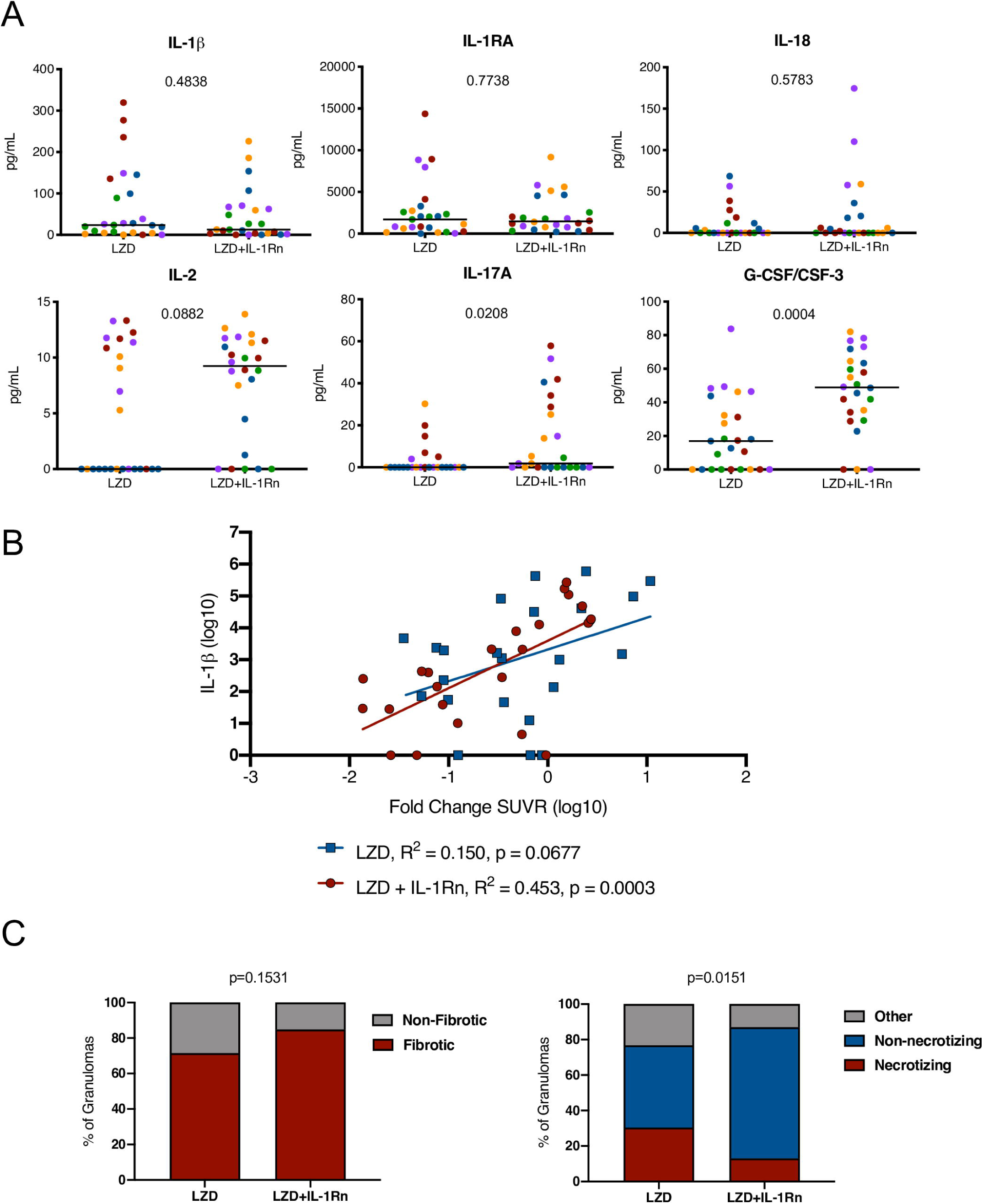
Inflammation modulation by HDT influences granuloma resolution. (A) Randomly selected granulomas (n=5 per macaque, n=25 per treatment group) were subjected to multiplex cytokine analysis to determine differences between treatment groups (colors indicate lesions from a single animal). p values shown determined by Mann-Whitney test. (B) A linear regression analysis of IL-1β levels and fold change in SUVR is depicted for LZD (blue) and LZD+IL-1Rn (red) granulomas. R^2^ and p values are provided in the legend. (C) Granulomas from LZD (n=57) and LZD+IL-1Rn (n=45) were examined by a pathologist (EK) and categorized. We compared fibrotic (red) vs non-fibrotic (gray) (left) frequencies between treatment groups. The right compares necrotizing (red) vs non-necrotizing (blue) and other (gray) granuloma frequencies. Fisher’s exact test p-values are reported.

## Discussion

HDTs are a tantalizing solution to improve TB therapy, however the complexities of host-pathogen interactions and host variability call for rigorous pre-clinical testing before implementation in humans. Here, we sought to determine the efficacy and safety of an HDT for TB comprised of LZD and IL-1Rn. IL-1 plays a complex role during TB, as IL-1 is important for early control of infection (7, 8, 34–36), yet is also associated with pathology at later times(10, 11). To validate the safety of IL-1Rn in this context, we first assessed the effects of IL-1 inhibition on TB disease progression in mice, using treatment regimens designed to concentrate on established disease. In C57BL/6, Nos2^-/-^ and C3HeB/FeJ mice, αIL-1R1 blockade reduced PMN infiltrates that are associated with pathological inflammation and did not impair bacterial control. The effect on bacterial control differs from a recent report in which lung bacterial burdens were increased by a similar αIL-1R1 treatment in C57BL/6 or C57BL/6 animals carrying the super susceptibility to tuberculosis-1 (Sst1) allele(25). We hypothesize that these differences are related to the timing of treatment, relative to the onset of adaptive immunity. Once infection is established, bacterial control is mediated predominantly via T cell functions, potentially reducing the importance of IL-1 dependent mechanisms(37). According to this model, the effect of IL-1 inhibition will depend on the timing of administration, as well as experimental factors that alter the timing of T cell priming and expansion, such as bacterial strain and dose. Regardless, our data indicate that IL-1 blockade can be beneficial in the context of established disease.

Any HDT will be administered in conjunction with antimicrobial therapy. When co-administered with LZD, blockade of IL-1R1 reduced PMN infiltrates in the mouse models, compared to the antibiotic alone. In macaques, IL-1Rn in conjunction with LZD significantly reduced overall lung inflammation as assessed by PET CT while LZD alone was more variable. IL-1Rn did not significantly enhance bacterial clearance compared to LZD alone, which was not surprising as we did not expect IL-1Rn to have direct antibacterial functions. However, it was important to assess bacterial burden during IL-1Rn treatment to ensure IL-1 blockade did not modulate host-mediated antibacterial mechanisms in granulomas and lymph nodes. In CFU positive lymph nodes, we observed a slight reduction in bacterial burden with IL-1Rn that approached significance (p=0.0965). Further studies should include assessing the influence of drug treatment and immunomodulation in lymph nodes, as thoracic lymph nodes are recognized as a site of Mtb persistence (29)

To further assess the effects of IL-1Rn modulation of immune responses, we observed a decrease in neutrophils (PMNs) and a corresponding decrease in IL-8 in the airways of both treatment group (LZD or LZD+IL-1Rn). This indicated LZD alone was efficacious in reducing PMN inflammatory signatures, likely due to the reduction in bacterial burden. In granulomas, levels of IL-1β, IL-1RA and IL-18 were not affected by IL-1Rn treatment in the subset examined. As IL-1Rn does not affect inflammasome formation and function it is not surprising that IL-1β and IL-18 production were minimally modulated by IL-1Rn. IL-1RA levels remained unchanged between treatment groups, which could be due to our method of detection or kinetics of consumption. Our multiplex analysis is specific for non-human primates and may not recognize recombinant human IL-1RA like Anakinra. In LZD+IL-1Rn treated granulomas we found a strong positive correlation between IL-1β levels and fold-change in SUVR, a surrogate for inflammation. These data highlight the relationship between IL-1β and inflammation in granulomas, further supporting the potential of IL-1 blockade as an HDT. In TB, IL-2 and IL-17A have protective functions however are primarily expressed by T cells (31). While IL-1β has been reported to enhance IL-17A production by T cells in mice, these studies have primarily used model-antigen systems and may not reflect a chronic infection like TB (38). IL-1 exacerbates pathology later in TB infection, indicating the effects of IL-1 on adaptive immunity are complex and likely dependent on the host system and infection model being used. G-CSF was increased in granulomas from LZD+IL-1Rn treated macaques which, combined with our observations of reduced PMN infiltrates in mouse lungs and macaque airways, indicates a paradoxical role of G-CSF as a neutrophil differentiation factor (39). However, in a model of LPS-induced lung injury G-CSF blockade induced accumulation of PMNs and increased inflammation in the lungs, indicating pulmonary inflammation may not follow dogmatic rules of canonical inflammatory pathways (39).

Lung fibrosis is modulated by the IL-1 pathway and while fibrosis is associated with healing tissue, lung fibrosis can cause secondary complications after TB disease resolution (2). We have shown in previous studies that TB drug therapy induces fibrosis in granulomas (32, 40). These studies in combination with the correlation between IL-1β and SUVR led us to assess whether LZD+IL-1Rn was associated with changes in granuloma pathology. While we did not observe a difference in fibrosis, there was a significant decrease in frequency of necrotizing granulomas with IL-1Rn in addition to LZD therapy. Thus, while there is no synergistic effect between IL-1Rn and LZD in promoting fibrosis associated with drug clearance, the reduction in neutrophils observed in mice and macaques could skew granuloma resolution towards a non-necrotizing lesion (41). Further studies with IL-1Rn alone could elucidate these dynamics, however we designed this macaque study with intention to be translatable to humans. We could not justify an IL-1Rn only treated group as TB-patients would receive anti-microbial therapy with any HDT or immunomodulatory intervention. However, our findings give precedence for further exploration of the potential of IL-1Rn as an HDT for TB and will require additional studies to determine mechanism and safety.

Our second goal was to abrogate LZD-associated bone marrow suppression with IL-1Rn therapy, thereby increasing the efficacy and therapy potential of LZD in TB. While we designed our HDT to match the current FDA guidelines for LZD and IL-1Rn schedules, this resulted in a lack of observable bone marrow suppression in macaques. In mice however, LZD-induced bone marrow suppression was reduced with the addition of IL-1R1 antagonist, supporting our initial hypothesis of the therapeutic potential of IL-1Rn in reducing LZD toxicity. The role of IL-1 signaling is consistent with the ability of IL-1 to suppress erythropoiesis in mice by reducing the number of progenitors (42). The remaining deficit in erythropoiesis during IL-1 blockade could reflect either incomplete inhibition by IL-1Rn or an independent role of inflammasome activation. Optimizing this effect will require further work to understand the relative roles of IL-1 signaling and inflammasome activation.

Overall, our data support previous findings in macaques that LZD has superb efficacy against TB, administered here as a single-drug given for only 4-weeks, supporting LZD as an antimicrobial for MDR/XDR-TB cases (12). IL-1Rn therapy in conjunction with LZD was successful in reducing TB-associated inflammation with no negative effects on bacterial clearance. The differences in bone marrow suppression between mice and macaques highlights the importance of assessing HDTs in multiple models prior to human trials, as there are clear biological differences between the two models. Anakinra (IL-1Rn) is already FDA-approved for adult and pediatric treatment of other inflammatory disorders and our data provide pre-clinical evidence for IL-1Rn consideration as potential therapy in TB. Further safety and mechanism assessments in translatable animal models are required, but one could envision IL-1Rn therapy to limit destructive inflammation for severe cases of TB. While our study emphasizes the importance of rigorous testing of HDTs for TB in multiple translational models prior to implementation in human trials, we also show that immunomodulation of the IL-1 pathway did not exacerbate TB disease.

## Supporting information

All Supplemental Figures, Tables and Legends

## Author Contributions Statement

CW, BM, CS and JF drafted and edited this manuscript. BM, JP, MS, and SN contributed to mouse experiments and/or figures. CW, PM, CC, CA, BS, HB, AW, EC, MZ, VD, PL and JF contributed to macaque experiments, data and statistically analysis and/or figures. All authors approved the final version of this manuscript.

## Acknowledgements

We thank all members of the Flynn, Sassetti, Mattila, and Lin laboratories for their collaborative efforts during this study. We are grateful for intellectual contributions from Dr. Clifton Barry III and Dr. Robert Wilkinson. Special thanks to all the veterinary and research staff for their efforts, expertise and dedication to these studies.

This manuscript has been released as a pre-print at bioRxiv (Winchell et al) (43).

## Data Availability Statement

All data are provided in a supplemental table.

