## Supplementary material for "Evaluation of IL-1 blockade as an adjunct to linezolid therapy for tuberculosis in mice and macaques": All Supplemental Figures, Tables and Legends

**Supplemental Data**

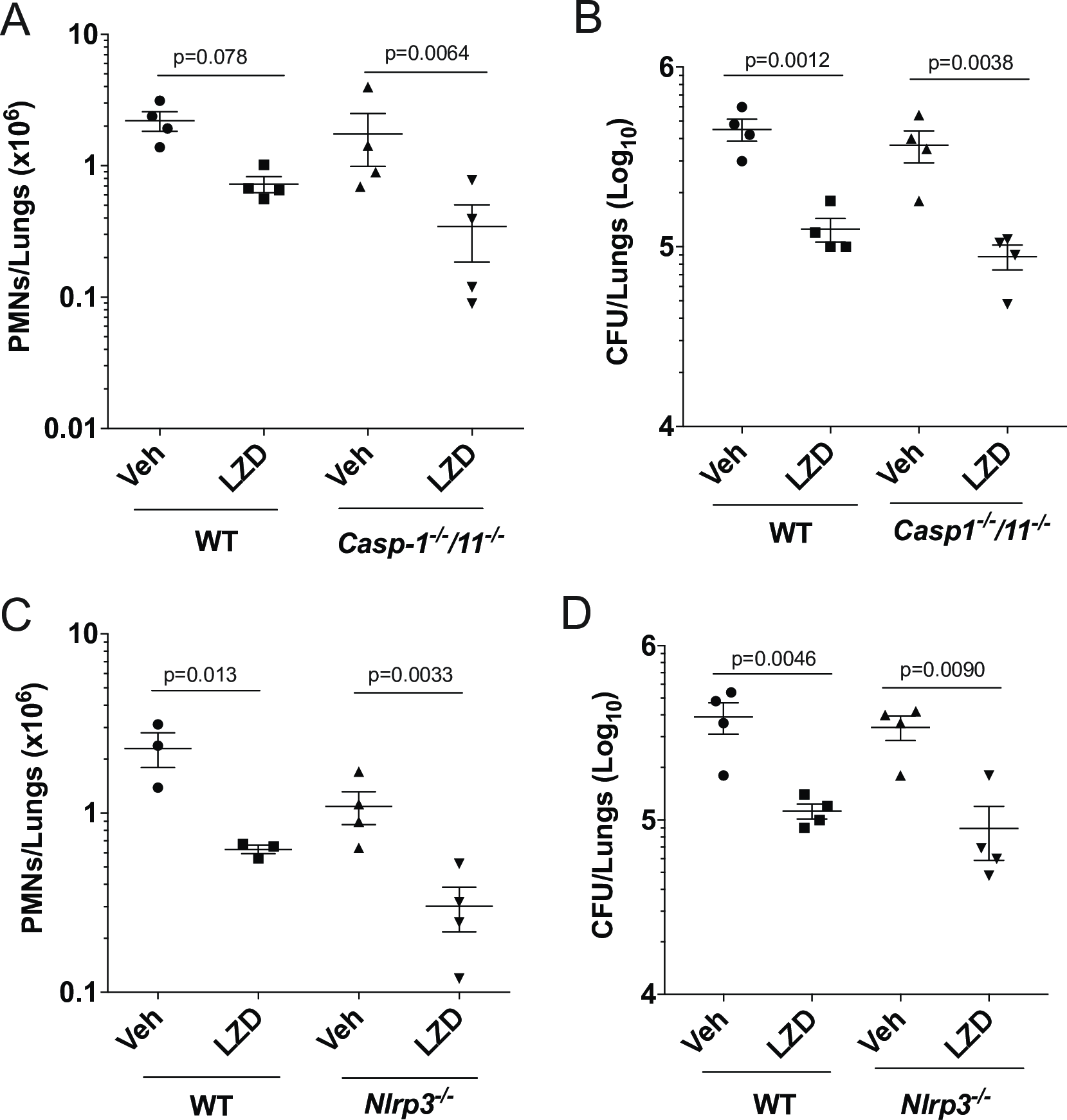

**S1 Fig. Inhibiting inflammasome signaling does not interfere with LZD efficacy.**

(A) Wild type C57BL/6 and *Caspase-1/11^-/-^* mice were infected with *M. tb* Erdman for 2 weeks, and treated with linezolid for the subsequent 2 weeks. (A) Lung neutrophil (PMN) infiltration was quantified by flow cytometry. (B) CFU in the lung are shown. Data shown (Mean ± SD) are representative of two independent experiments. (C) WT and *Nlrp3^-/-^* mice were infected with *M. tb* Erdman for 2 weeks and treated with linezolid for the following two weeks. (C) Lung neutrophils were quantified by flow cytometry. (D) Bacterial burden in the lung were quantified as CFU. Data shown (Mean ± SD) are from one experiment. One-way ANOVA with Sidak’s multiple comparison test was used to calculate the indicated p-values. N=3-5 mice used for each treatment group.

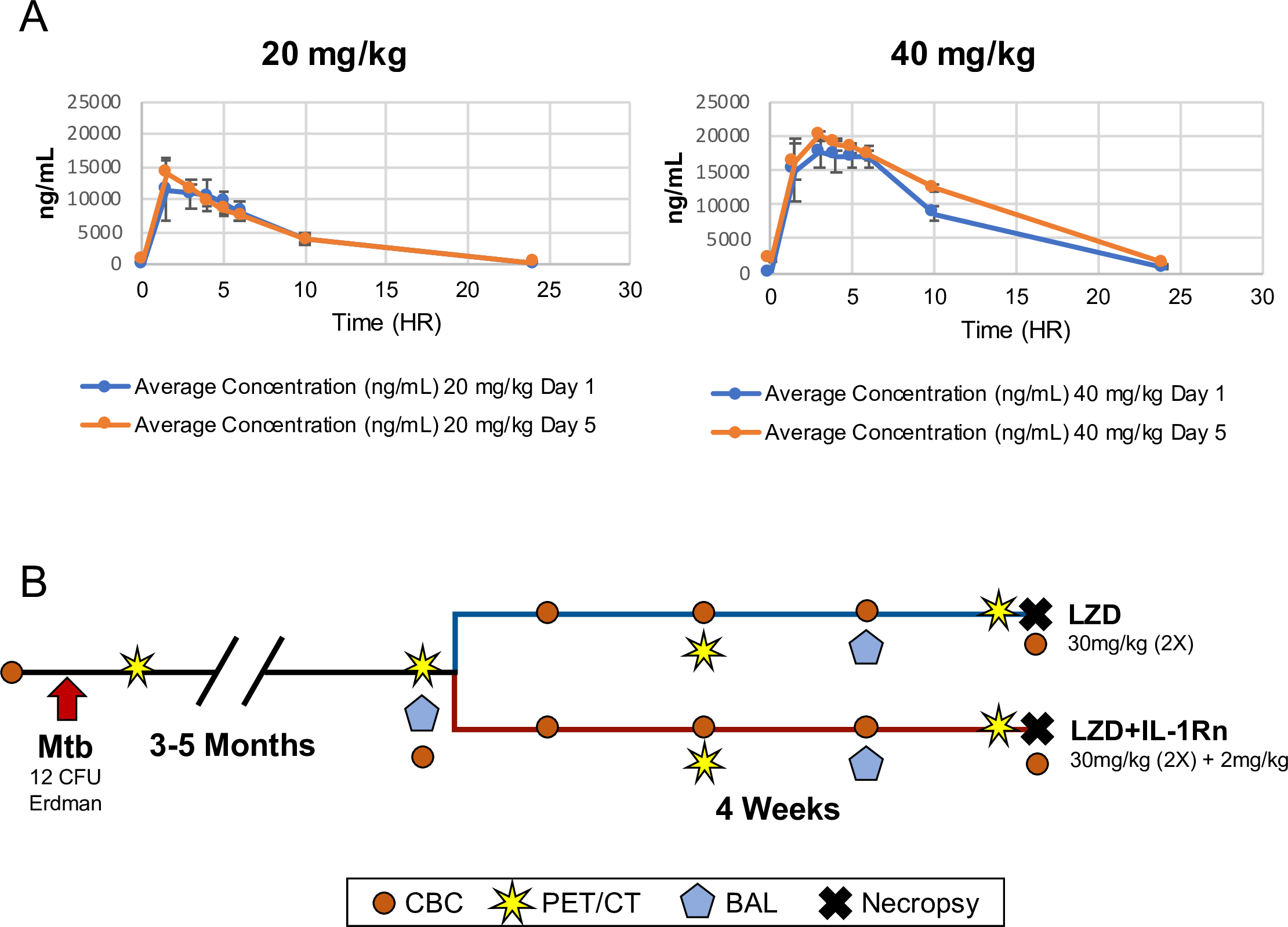

**S2 Fig. Linezolid pharmacokinetic analyses and Experimental plan for HDT in macaques**

(A) Plasma concentration time profiles of cynomolgus macaques receiving 20mg/kg or 40mg/kg linezolid. Average plasma concentrations were plotted on Day 1 and Day 5 post-dose (n=3) (B) Experimental timeline of macaque study.

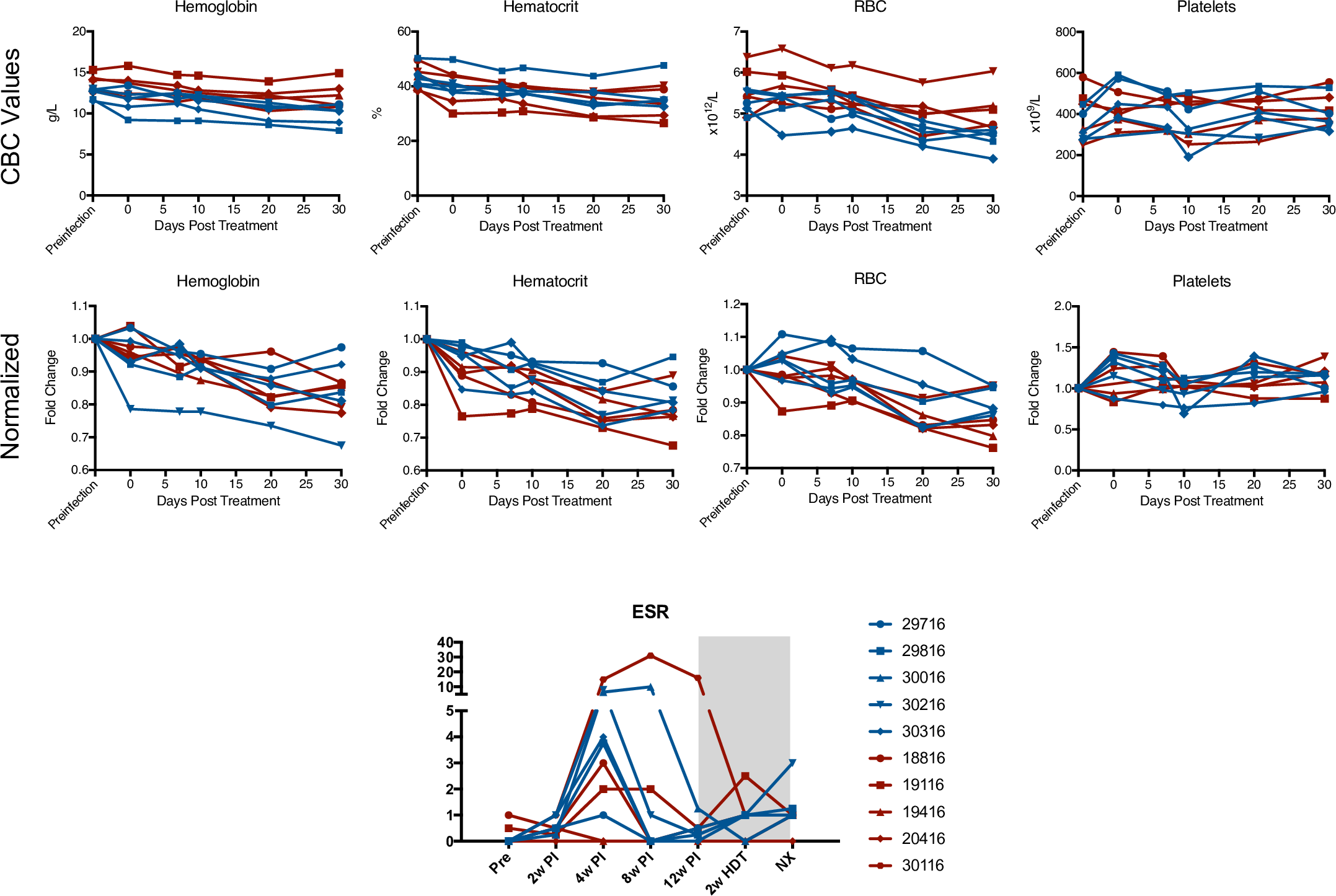

**S3 Fig. Complete blood counts and erythrocyte sedimentation rate for macaques undergoing HDT**

Complete blood counts (CBCs) were taken through the study to assess anemia caused by linezolid or Mtb infection. The CBC values (top) show true values for each macaque in LZD (blue) or LZD+IL-1Rn (red) treatment groups. Due to variability in animal size and age, we normalized CBC values to the preinfection time point for each animal (middle graphs). These data indicate the relative change over the course of HDT. ESR (bottom) is a non-specific measure of inflammation, LZD (blue) and LZD+IL-1Rn (red) treated macaques are depicted.

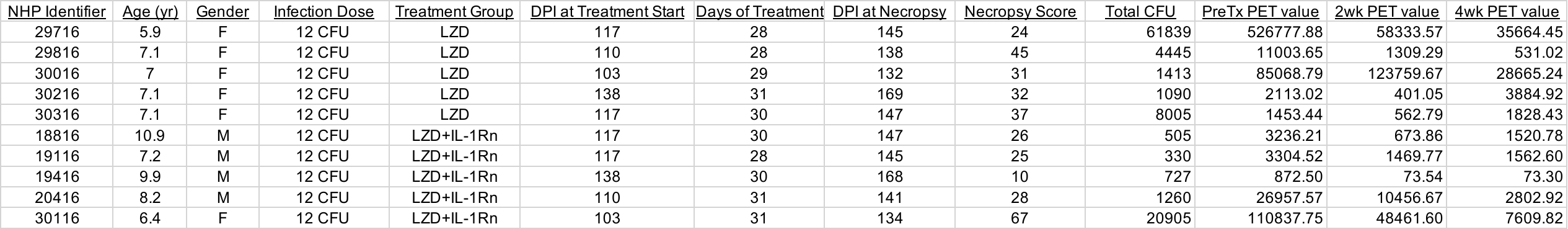

**Supplemental Table 1 - Macaque information and PET CT data.**
Provided are details regarding macaques utilized for this study. DPI = days post infection.
